# A Single Cell Atlas of Spared Tissue Below a Spinal Cord Injury Reveals Cellular Mechanisms of Repair

**DOI:** 10.1101/2021.04.28.441862

**Authors:** Kaya J.E. Matson, Daniel E. Russ, Claudia Kathe, Dragan Maric, Isabelle Hua, Jonathan Krynitsky, Randall Pursley, Anupama Sathyamurthy, Jordan W. Squair, Gregoire Courtine, Ariel J. Levine

## Abstract

After spinal cord injury (SCI), the “spared” tissue below the lesion contains undamaged cells that could support or augment recovery, but targeting these cells requires a clearer understanding of their injury responses and capacity for repair. Here, we used single nucleus sequencing to profile how each cell type in the lumbar spinal cord changes over time after a thoracic injury. We present an atlas of these dynamic responses and explore two unexpected findings. Amongst neurons, rare cell types expressed a molecular signature of regeneration and amongst microglia, we identified a population of “trauma associated microglia” (TAM). These TAM cells were present in the white matter near degenerating axons and expressed the trophic factors *Igf1* and *Spp1*(OPN). Viral over-expression of *Igf1* and *Spp1*(OPN) expanded the TAM population and promoted the clearance of myelin debris. These findings expose endogenous mechanisms of repair in spared neural tissue, uncovering potential candidates for targeted therapy.

## Introduction

The brain, spinal cord, and peripheral nervous system are comprised of diverse cell types that operate together as local communities to enable normal physiology. Following acute trauma, a complex interplay of cellular responses shapes the outcome. Whether the tissue can restrict the damage, promote structural remodeling and functional compensation, and ultimately achieve recovery depends on myriad dynamic molecular changes amongst neurons, astrocytes, microglia, oligodendrocytes, vascular cells, and many other cell types^1,2^.

Spinal cord injury (SCI) is a traumatic event that can cause long-lasting paralysis, pain, autonomic dysregulation, and body-wide physiological changes^3^. Understanding and developing therapeutics that target cellular changes within the lesion epicenter is undoubtedly valuable. In addition to the lesion of the SCI, there is an emerging focus on the “spared tissue” below a SCI and its inherent potential for supporting anatomical reorganization and recovery in people with SCI. For example, epidural electrical stimulation of the lumbar spinal cord, combined with rehabilitation training, can promote impressive gains in motor function and autonomic control and provide enhanced quality of life^4-6^. This therapy underscores the importance of understanding the spared tissue below the site of injury to reveal the intrinsic potential for spontaneous and rehabilitation-driven recovery.

Here, we sought to uncover the dynamic cell type specific responses of spared lumbar spinal cord following SCI with the goal of identifying the molecular and cellular mechanisms that promote or restrict recovery. First, we performed severe contusion spinal cord injuries in mice and tracked the progression of injury responses from acute to chronic timepoints. To profile the diverse cell types within the lumbar spinal cord following thoracic injury, we used single nucleus RNA Sequencing (snRNA-Seq) and created an “atlas” of the spared tissue after injury. The size and scope of this dataset allowed us to identify rare cellular populations that displayed molecular pathways with the potential to support recovery. We identified neuronal populations that induce a transcriptional signature of regeneration, providing a template for expanding this potential in the future. Finally, we identified a population of “trauma associated microglia” that were localized along white matter tracts of degenerating axons and that used autocrine signaling via IGF1 and osteopontin (*Spp1*/OPN) to promote proliferation. This population could be expanded with exogenous IGF1 and *Spp1*/OPN to degrade, phagocytose, and clear local myelin. Together these findings shed light on the limited spontaneous mechanisms of repair in the spared tissue below a SCI but also reveal latent potential for targeted neuro-regeneration and tissue remodeling therapies.

## Results

### Creating a Single Cell Atlas of the Spared Lumbar Cord After Spinal Cord Injury

We sought to understand how the spared tissue below a spinal cord contusion responds to the injury. A severe contusion was delivered to the thoracic spinal cord of mice, resulting in a near complete loss of neurons at the site of injury (Extended Data Fig 1) and immediate paralysis (Fig. 1a). Locomotor function was tracked over a range of timepoints that, in mice, corresponded to the acute injury period (1 day post injury (dpi)), the sub-acute and intermediate stages that are typically targeted with therapeutic interventions (1 week and 3 weeks post-injury (wpi)), and a chronic time at which point recovery typically plateaus (6 wpi; Fig. 1a). We then dissected the entire lumbar spinal cords of animals from each condition in triplicate and performed single nucleus isolation and RNA sequencing (Fig. 1b)^7^. After integration and the removal of low-quality nuclei and doublets, we obtained a total set of 67,903 nuclei.

**Figure 1:**
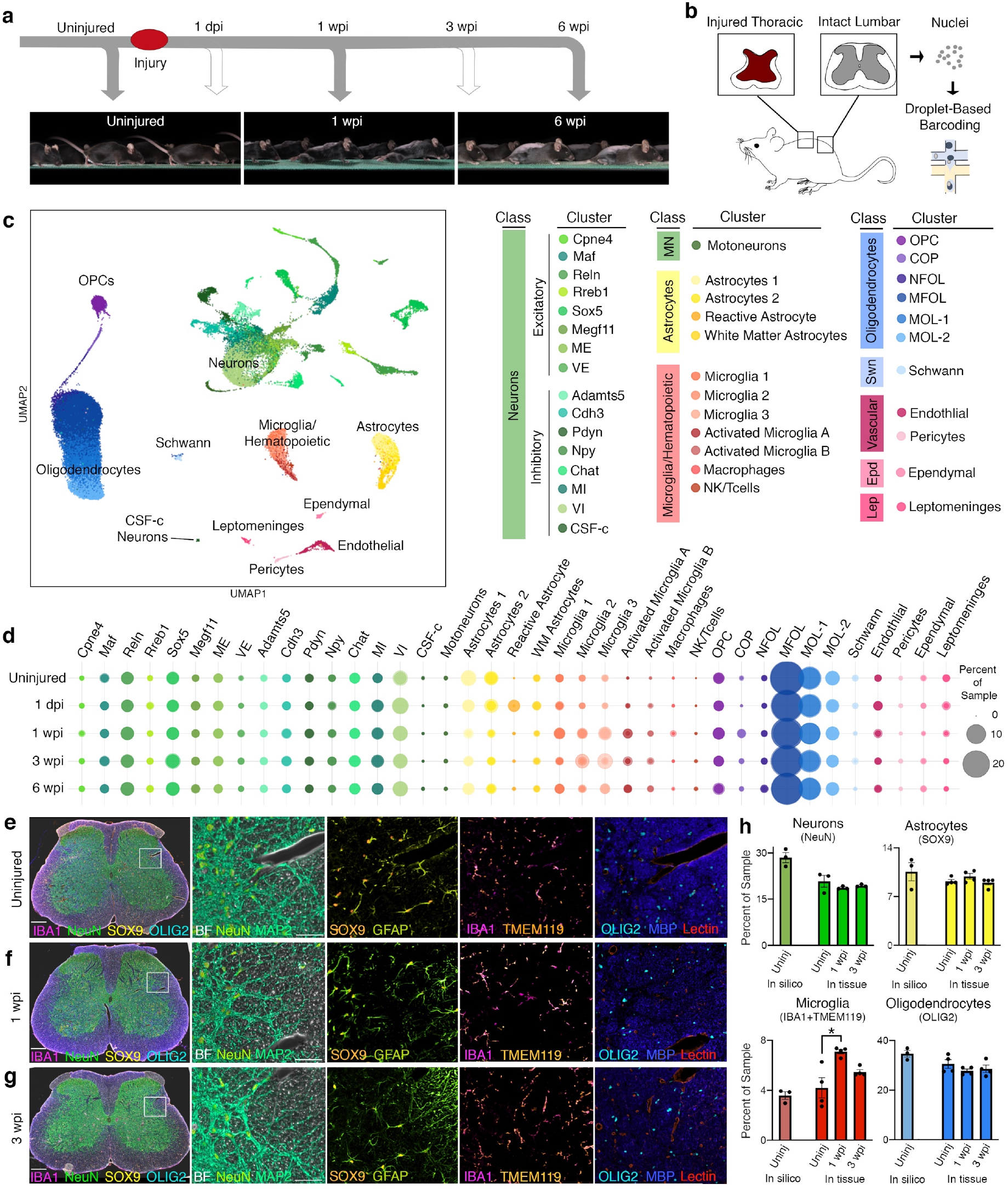
Single Nucleus RNA Sequencing of the Lumbar Spinal Cord after Thoracic Contusion. **a**. Top, an overview of experimental design for injury and tissue collection. Bottom, example chronophotography from uninjured, 1 wpi and 6 wpi mice. **b**. Schematics depicting the experimental design for snRNAseq, showing the injured thoracic cord and intact lumbar cord (with dark red representing the lesion) as well as nuclei isolation from the intact lumbar cord followed by droplet-based barcoding for single nucleus RNA sequencing. **c**. Uniform manifold approximation and projection (UMAP) visualization of 67,903 nuclei from uninjured and injured lumbar spinal cords, revealing 8 classes and 39 subtypes. **d**. A bubble plot representing proportion of a sample from a given cell type (N = 3). **e-g**. Multiplex Immunohistochemistry of the lumbar spinal cord from uninjured, 1 wpi and 3 wpi. Tissue was stained for NeuN (lime green), MAP2 (seafoam green), Sox9 (orange), GFAP (yellow), IBA1 (magenta), TMEM119 (peach), OLIG2 (cyan), MBP (blue), Lectin (red), and brightfield was imaged. Scale bars are 200 µm and 50 µm, respectively. **h**. Quantification of the in silico and in tissue percentage of four cell classes. For in silico quantification of neurons and astrocytes, the proportion of total neurons was calculated relative to the sample. For microglia, the proportion of both activated microglia clusters and three homeostatic microglia clusters were used to calculate the proportion. For oligodendrocytes, OPCs, COPs, NFOL and MFOL were used to calculate the proportion (of OLIG2-expressing cells). In tissue percentages were calculated using the percent of DAPI+ neurons (NeuN), astrocytes (SOX9), microglia (IBA1+TMEM119) or oligodendrocytes (OLIG2). Error bars indicate mean ± SEM (N = 3, in silico; N = 4, in tissue).

To create a single cell atlas of cell type specific responses to injury, we performed clustering of these nuclei in multiple hierarchical phases (see Methods). At a coarse level of clustering, the 8 major cell classes of the spinal cord are neurons, astrocytes, microglia/hematopoietic cells, oligodendrocyte lineage cells, Schwann cells, endothelial cells, pericytes, ependymal cells, and leptomeninges (Fig. 1c, Extended Data Fig. 2a). Each of these major classes were identified using well-established markers (Extended Data Fig. 2b). Neurons expressed *Snhg11, Rbfox1, Rbfox2, Snap25*, and were characterized either by the excitatory marker *Slc17a6*, the inhibitory markers *Gad1* and *Gad2*, or the cholinergic markers *Slc5a7* and *Chat*. Astrocytes expressed *Slc7a10, Agt, Gfap*, and *Vim*. Microglia/hematopoietic cells expressed *C1qa, Ctss, Gpnmb, Lgals3, Itgax, Ms4a4b, Cd3g* and *Nkg7*. Oligodendrocyte lineage cells including oligodendrocyte precursor cells (OPCs) expressed *Cspg5* and *Tnr*; committed oligodendrocyte progenitor cells (COPs) expressed *Fyn* and *Tcf7l2*, whereas myelinating and mature oligodendrocytes expressed *Plp1, Mag* and *Mog*. Schwann cells expressed *Mpz* and *Pmp22*. Vascular cells included endothelial cells, which express *Bsg* and *Cldn5* and pericytes, which express *Vtn* and *Pdgfrb*. Ependymal cells express *Nnat* and *Dnah12*. Leptomeninges express *Dcn* and *Col1a1* (Extended Data Fig. 2b, Extended Data Table 1). At a more resolved level, we observed 39 distinct cell types (Fig. 1c, Extended Data Fig. 2b). With this approach, we have identified the major classes and specific cell types of healthy and injured mouse spinal cord.

### Cell Type Compositional Changes in the Lumbar Cord After Injury

We next aimed to characterize how the cellular composition of the lumbar spinal cord changed after thoracic injury. To normalize for variable numbers of nuclei from each sample, the proportion of each cell type was calculated. In the uninjured tissue, oligodendrocytes represented largest proportion of the spinal cord, at 30.8% myelinating and 17.7% mature oligodendrocytes (average of three replicates, ± 0.8 and 0.8% SEM). Neurons were 27.0% of the uninjured spinal cord, astrocytes were 10.6% and microglia/hematopoietic cells were 3.7% (± 4.6, 1.6, 1.3%; Fig. 1d, Extended Data Table 2).

We found that the overall cellular composition stayed relatively stable after injury, with two notable exceptions that occurred within the first week following thoracic contusion. The proportion of microglia/hematopoietic cells tripled, while the proportion of astrocytes decreased to one third of uninjured levels over the same time course (p val < 0.05, Fig. 1d, Extended Data Fig. 6). In addition, there were relatively small changes in sub-types of the oligodendrocyte lineage, including a proportional decrease in myelinating oligodendrocytes and an expansion of committed oligodendrocyte precursors (Fig. 1d, Extended Data Fig. 4).

To determine whether these observations accurately reflect endogenous changes in the spared lumbar spinal cord, we performed immunohistochemical staining and in situ hybridization experiments in tissue sections from healthy animals or following one or three weeks after thoracic contusion injury. First, a highly multiplexed protein and cell stain assay was performed using multiple markers for each major cell class.^8^ This method complements the highly dimensional single cell sequencing approach of cell type identification with the richness of spatial information. We stained for neurons (NeuN, MAP2, Neurofilament-L, Neurofilament-M and Neurofilament-H), astrocytes (GFAP and SOX9), oligodendrocytes (CNPase, MBP, and OLIG2), neuroinflammatory cells (TMEM119, IBA1, CD68, CD11c), and vasculature (isolectin B4, tomato lectin, smooth muscle actin). Quantitative analysis of the multiplexed immunohistochemistry validated the stable proportions of neurons (20.8% uninjured ±0.6, 17.7% 1 wpi ±0.7, 18.5% 3 wpi ±0.7, ns) and oligodendrocytes (30.5% uninjured ±0.4, 27.7% 1 wpi ±0.3, 28.6% 3 wpi ±0.4, ns) as well as the significant increase in the number of microglia within the spared lumbar spinal cord (4.7% ±0.3 at uninjured, 8.5% ±0.2 at 1 wpi, respectively; Fig. 1e-h). In addition, the numbers of each cell type in tissue were found to be similar to the proportions observed in the sequencing analysis, supporting the validity of these independent approaches.

This in-tissue analysis did not confirm a major decrease in the proportion of astrocytes amongst spinal cord cell types following SCI that was observed in the sequencing data. Rather, we observed no significant change in SOX9-expressing astrocytes (Fig. 1h). To further characterize the proportion of astrocytes using combinatorial RNA expression that would be more similar to the single nucleus sequencing data, we performed multiplexed RNA in situ hybridization using *Agt, Gja1*, and *Aqp4* markers. In this context, a modest decrease was observed from 11.7% in uninjured lumbar cords to 9.5% 1 wpi (±0.7%, 0.9%, respectively, Extended Data Fig. 3c). This suggests that the proportional change in astrocytes that we observed in the single cell atlas did not reflect endogenous cell type changes but is likely due to the overall proportional shifts of cell types in the injured spinal cord.

**Figure 2:**
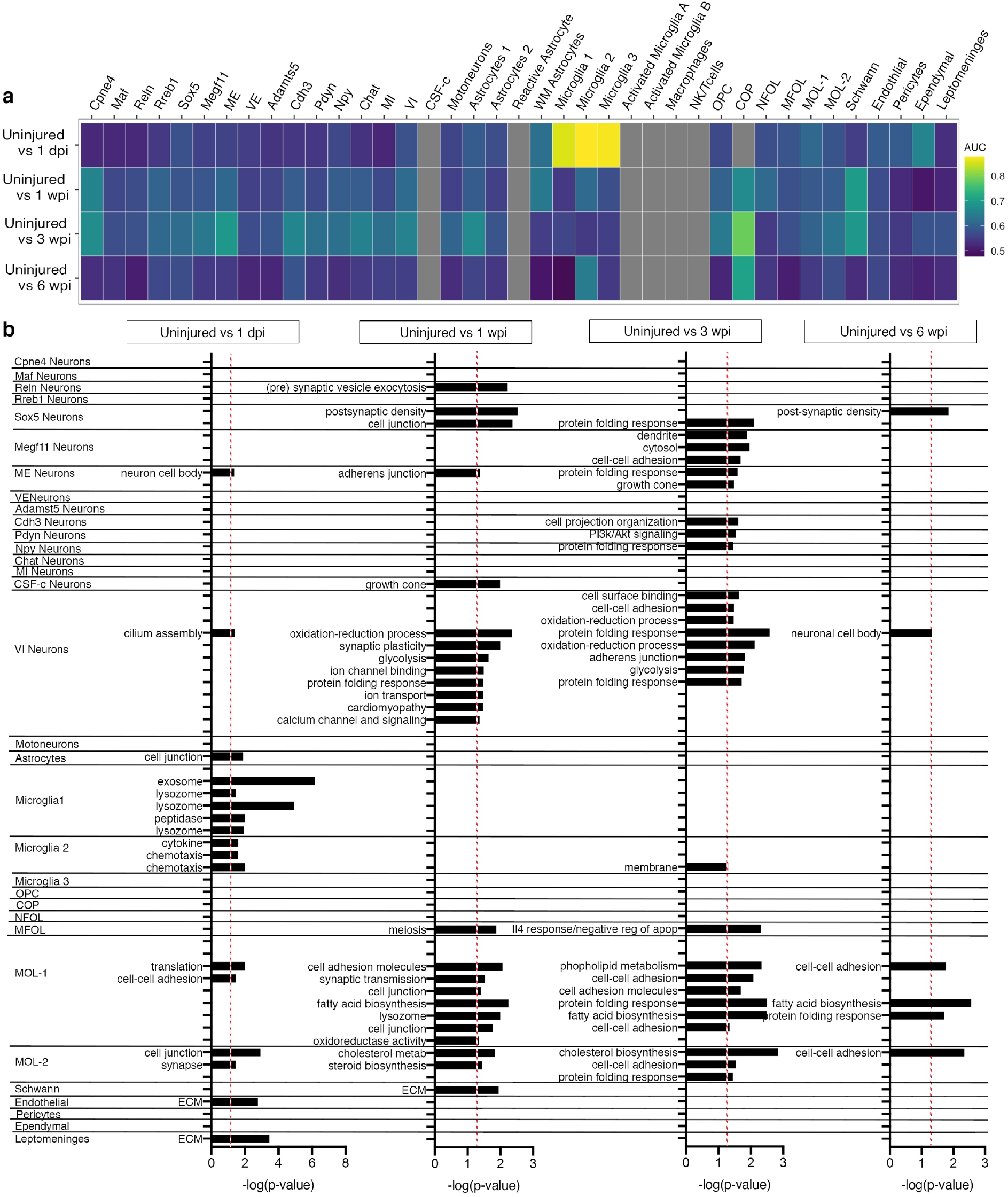
Cell Type Specific Changes in Gene Expression after Injury. **a**. Cell types ranked based on responsiveness by Augur. AUC score represented from dark blue to yellow. Clusters with insufficient number of cells in the uninjured timepoint are plotted in grey. **b**. Pathway analysis for differentially expressed genes between uninjured and injured timepoints. X-axis indicates −log(p value) of GO and KEGG pathway clusters. Red dotted line indicates −log(p-value) 1.3 (p value 0.05).

**Figure 3:**
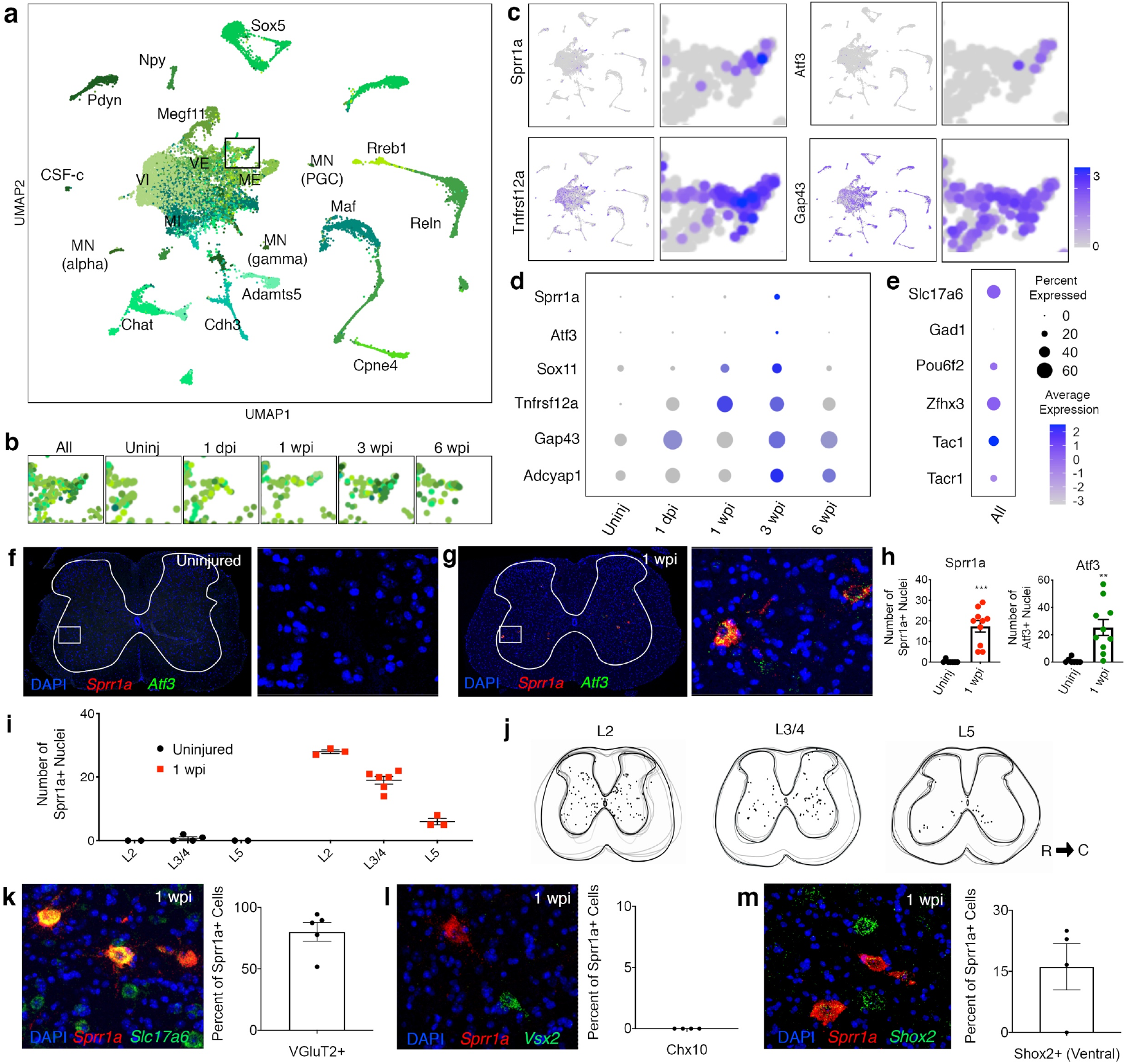
Specific Neurons Express Genes Associated with Regeneration. **a**. UMAP showing predicted neuronal families. **b**. Targeted view of RAG-expressing cluster over injury time points. **c**. Featureplots and violin plots showing RAGs expressed in neurons. **d**. A dotplot showing expression of RAGs within targeted RAG-expressing cluster (cluster 23 defined by independent clustering in Extended Data Fig. 5), split by timepoint. **e**. A dotplot showing marker genes for the cluster expressing RAGs **f-g**. RNAscope *in situ* expression of *Sprr1a* and *Atf3* in the uninjured cord and 1 wpi. **h**. Quantification of *Sprr1a*+ cells and *Atf3*+ cells in the uninjured and 1 wpi injury cord (N = 7, 10). **i**. Rostral-caudal quantification of *Sprr1a*+ cells in the spinal cord (N = 4, 6). **j**. A heatmap representation of *Sprr1a*+ cells (N = 3, 6, 3 animals). **k-m**. Visualization and quantification of VGluT2/*Slc17a6*+, Chx10/*Vsx2*+, and *Shox2*+ *Sprr1a*-expressing cells in the ventral spinal cord. Error bars indicate ± SEM (N = 5, 4, 4 animals).

Overall, we found that the cellular composition of the spared lumbar spinal cord does not change considerably after SCI. Against this stable background, there was a pronounced increase in microglia, though the lumbar cord that we profiled is millimeters away from the injury site. We also emphasize the importance of in situ validation for cell proportion changes in single cell data, as it is challenging to distinguish relative changes in cell type balance from authentic changes in endogenous cell proportions.

### Cell Type Specific Changes in Gene Expression in the Lumbar Spinal Cord After Injury

While the overall balance of cell types is relatively stable after injury, we hypothesized that gene expression *within* cell types would change after injury, endowing new cell type properties and functions. Our goal was to determine which cell types changed significantly following injury. However, it is important to note that statistical differences in gene expression can largely be driven by the size of the cluster, with larger clusters having the power to resolve more differentially expressed genes. We therefore implemented Augur, a method to rank responsiveness of cell types in single cell data that is not biased by cluster size^9^. We applied Augur to the 39 cell types, including 17 neuronal clusters, 4 astrocyte clusters, 7 microglial/hematopoietic clusters, 6 oligodendrocyte related clusters, Schwann cells, endothelial cells, pericytes, ependymal cells, and leptomeninges. We found that the average cell type prioritization score across all cell types decreased with time post-injury, suggesting a return to homeostatic conditions (average 0.55 AUC, Fig. 2a). At 1dpi we found that microglia were the most perturbation-responsive cell type (0.86 average AUC for Microglia 1, 2, and 3). These results suggest that microglia play an important role in the immediate phase after injury, even in spared tissue distant from the site of injury.

To understand the cellular processes driving these changes, we examined differentially-expressed genes within these cell types after injury and performed gene ontology (GO) and pathway analysis. At 1 dpi, glial cells changed the most. Microglia/hematopoietic cells changed dramatically and were characterized by a burst of expression of genes related to cell metabolism, which may support the significant expansion in microglial cell numbers that we observed above. Both microglia/hematopoietic cells and astrocytes displayed signs of activation despite their considerable distance from the lesion site in the thoracic cord. Reactive astrocytes emerged 1 dpi, expressing pan-reactive genes such as *Lcn2* and *Serpina3n*, as well as markers for pro-inflammatory and neuroprotective astrocytes^10^ such as *Gbp2* and *Slc10a6* (Extended Data Fig. 3). Other glial cell types such as endothelial cells and meninges altering molecules related to structure, cell-cell connections, and extra-cellular matrix (ECM) adhesion. In addition, nearly all cell types showed increased expression of *Mt1* (metallothionein-1) and *Fth1* (Ferritin Heavy Chain 1) genes which are both involved in binding heavy metals including iron. This suggests that extravascular blood may be an early environmental cue to reach the spared tissue of the lumbar cord (Fig. 2b). Overall, the largest changes in gene expression occurred 1 dpi.

One week post injury, neuronal populations began to change by altering the expression of genes related to cell stress including oxidation-reduction processes and the protein folding stress response, as well as molecules related to neurotransmission and ion channel activity. Intriguingly, specific populations of excitatory neurons in the dorsal horn and inhibitory neurons within the ventral horn displayed changes in genes related to synaptic organization and plasticity. This suggests the potential for tailored circuit remodeling. At the same time, oligodendrocytes altered their cellular metabolism molecules and genes related to cell-cell adhesion. Three weeks after injury, multiple neuronal populations continued to show signatures of cell stress and changes in cell-cell contacts, while oligodendrocytes continued to show changes related to cell metabolism and cell-cell adhesion. Finally, by 6 wpi, many cell types showed modest or no changes in gene expression compared to the uninjured state, mirroring the Augur findings above which showed an overall return to baseline expression patterns by this point (Fig. 2b).

Together with the cellular composition analysis above, these data comprise a natural history time course of how the lumbar spinal cord responds to injury. Within each time window or epoch, the community of cell types responds with diverse molecular signatures and within each cell type, a specific molecular trajectory can be observed across time.

### Rare Populations of Spinal Neurons Induce a Gene Expression Signature of Regeneration

Neurons in the spared lumbar spinal cord spinal are thought to mediate partial recovery in multiple therapeutic contexts, though the molecular mechanisms that mediate this function are mostly unknown. We observed broad but modest changes in the expression of genes related to neural excitability, plasticity, and circuit structure that could support tissue-wide changes in function (Fig. 2b). But we also sought to explore whether, upon this background, there were also specialized adaptations to injury within highly refined populations of spinal neurons. Such changes would be challenging to observe without the resolution of single nucleus sequencing and thus the dataset that we generated held special opportunities for discovery.

We determined the refined identities of neuronal populations in the dataset by comparison with the neuronal populations described in a recent meta-analysis of mouse spinal cord cell types^11^. Label transfer was used to annotate neuron identities in an automated manner at the resolution level of “families”^12^. While we observed nearly all of the cell types that we expected, one group of neurons remained challenging to categorize (Fig. 3a, Extended Data Fig. 5a). Moreover, independent clustering of the neurons in this dataset suggested that this population represented a cluster with an unexpected expression profile that included genes associated with axon regeneration (Extended Data Fig. 5b-c).

“Regeneration associated genes” (RAGs) are induced in the peripheral nervous system after nerve injury, enabling regeneration of axons^13^, but most adult central nervous system neurons do not express these genes and lack the cell-intrinsic ability to substantially regenerate axons^14^. In this RAG-expressing cluster, we observed *Atf3, Sprr1a, Tnfrsf12a* (Fn14), *Sox11, Bdnf, Adcyap1*, and enriched expression of *Gap43* (Fig. 3c-d). Inspection of the other genes in this cluster suggested a mixed cell type origin. Cells with the highest RAG expression largely downregulated their endogenous gene expression, as has been previously reported in peripheral nerves that express RAGs, confounding molecular definition of their cell type^15,16^. We noted that this cluster expressed a diverse set of genes associated with ascending projection neurons from the lumbar spinal cord to the brain, including *Lypd1, Tacr1, Zfhx3, Pou6f2, Tac1, Syt4, Fam19a2, Scn9a, Nms*, and *Pde8b* (Fig. 3e)^17-21^. In particular, the expression of *Slc17a6* (vGlut2), *Zfhx3*, and *Pou6f2* suggested that some of these cells were likely excitatory neurons that resided in the lateral part of deep dorsal or ventral horn^21-23^. Together, these data suggest that a heterogeneous but rare set of ascending spinal cord neurons induce a RAG signature after contusion injury.

To determine whether spinal cord neurons induced the RAG signature in vivo, we analyzed the expression of *Sprr1a* and *Atf3* in the lumbar spinal cord and found rare RAG-expressing neurons starting at 1 wpi (Fig. 3f-h) through 6 wpi (Extended Data Fig. 5). A spatial analysis revealed a rostral-caudal gradient in the number of RAG-expressing cells, with more Sprr1a-positive cells at rostral lumbar segments closer to the lesion (Fig. 3i-j). The cellular distribution within the transverse plane also shifted such that RAG-expressing cells were found in the dorsal, mid, and ventral horns at L2 but were restricted to the ventral horn in L5. Based on the characteristic locations of these neurons, we hypothesized that they reflect post-synaptic dorsal column neurons and deep dorsal horn anterolateral tract neurons at L2/L3 and spinocerebellar neurons at L2 through L5. To test whether the ventral RAG-expressing neurons are excitatory, putative spinocerebellar neurons, we used a recently identified marker, *Shox2*, as well as the excitatory marker *Slc17a6*^24-26^. Indeed, we found RAG-expressing ventral neurons are nearly all excitatory and many express the *Shox2* marker as well (Fig. 3k-m). In addition to serving as a marker of spinocerebellar neurons, *Shox2* is also found in excitatory, rhythm-generating central pattern generator neurons of the ventral horn where they partially overlap with *Vsx2* (Chx10). Triple labelling for RAG markers, *Shox2*, and *Vsx2* demonstrated that only *Shox2* expressing neurons induce the RAG signature, revealing a high degree of specificity of this rare and unexpected pro-regenerative response (Fig. 3l-m).

Together, these data revealed that specific populations of spinal cord neurons induce a regeneration associated gene expression signature and that these are likely subsets of ascending projection neurons. It is possible that the ventral populations of neurons that induced RAG expression who’s axons run past the lesion site in the ventral white matter were injured but not axotomized by the contusion injury (which is delivered to the dorsal surface of the cord), while dorsal neurons that are more directly damaged do not induce a regenerative signature in this injury context. To determine whether complete axotomy elicits a distinct response, we performed complete transection injuries of the thoracic spinal cord and analyzed RAG expression in the lumbar spinal cord. One- and three-weeks following injury, we found a similar spatial pattern of RAG expression as was observed in contused spinal cords (Extended Data Fig. 5). Although most central nervous system neurons do not typically have an intrinsic capacity to regenerate, this raises the intriguing possibility that particular populations of spinal neurons may possess regenerative potential^14^.

### A Population of “Trauma Associated Microglia”

The compositional and gene expression data presented here revealed that microglia are the most perturbation-responsive cell type in the spared lumbar tissue after SCI (Fig. 1-2). To explore microglia and related hematopoietic cell types in greater depth, we independently clustered this class of cells and observed three homeostatic microglia populations, two activated microglia populations, a cluster of macrophages and a cluster of natural killer and T cells (Fig. 4a). Homeostatic microglial clusters were defined by *Cst3, C1qa, Ctss, Hexb, Trem2, P2ry12* and *Tmem119*. The activated microglia populations expressed lower levels of *P2ry12* and *Tmem119* and induced expression of the phagocytic markers *Cd68* and *Lyz2*. In addition, “activated microglia A” expressed *Gpnmb, Apoe, Lgals3, Igf1*, and *Spp1*, while “activated microglia B” expressed genes associated with pro-inflammatory microglia, such as *Ccl2, Ccl3, Ccl4* and *Lpl*. Macrophages expressed the genes *Mrc1, Cd74*, and *H2-Ab1* and did not express the microglia-specific genes *Tmem119* or *P2ry12*. Natural killer and T cells clustered together and expressed the genes *Ms4a4b, Cd52, Ptprc, Nkg7*, and *Cd3g*. All microglia/hematopoietic cell types increased in proportion relative to other cell types at 1 wpi (Extended Data Fig. 5). Both of the activated microglia sub-types were still present at six weeks post injury, suggesting that they may play an ongoing role in spared tissue (Extended Data Fig. 5).

**Figure 4:**
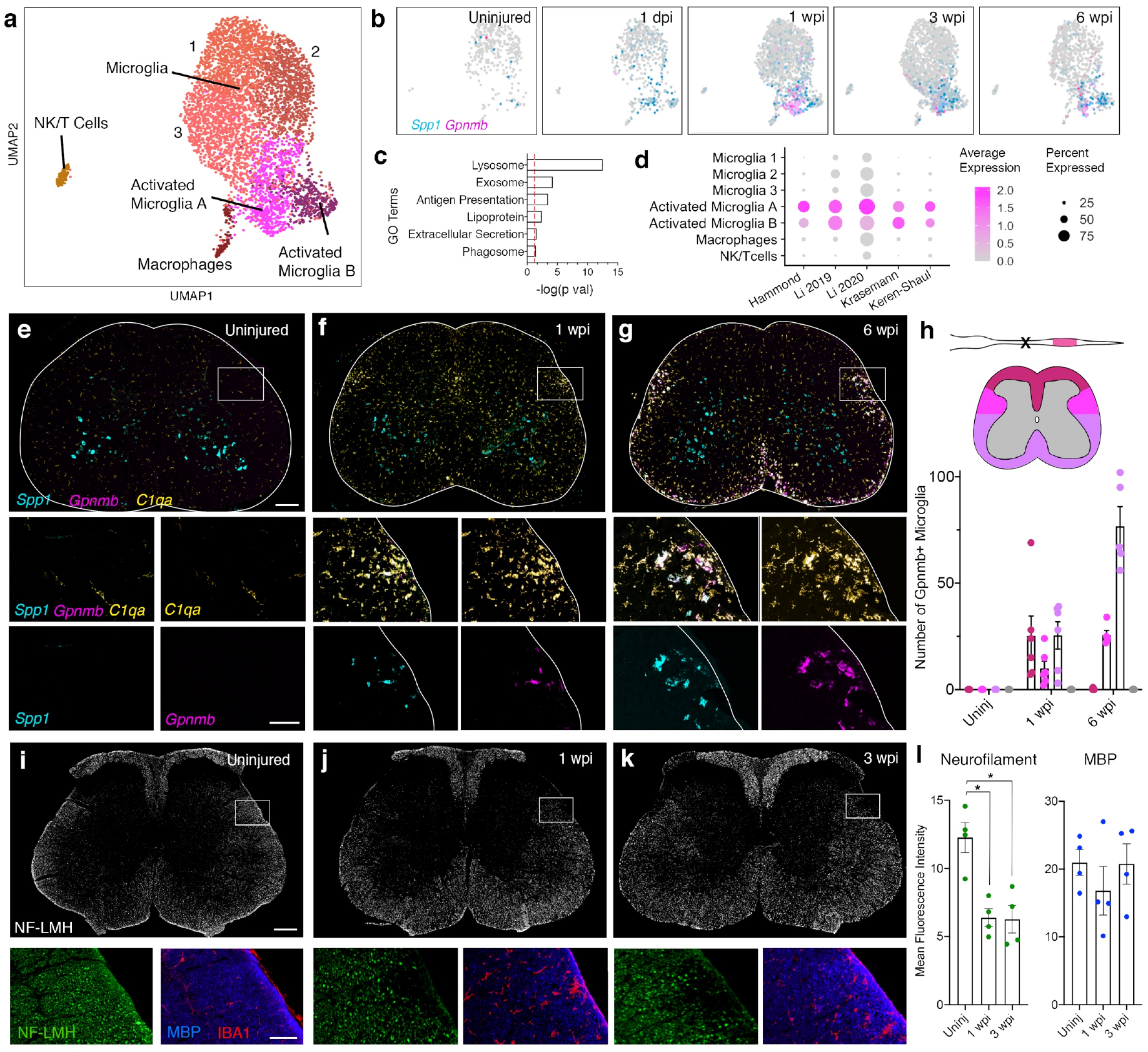
Microglia Expansion and after Injury and the Emergence of a “Trauma Associated Microglia”. **a**. UMAP of the 7 microglia subtypes. **b**. Featureplot of two marker genes of the “activated microglia A” population, *Spp1* (cyan) and *Gpnmb* (magenta), split by the timepoint (Uninjured, 1 dpi, 1 wpi, 3 wpi, and 6 wpi). **c**. GO analysis for “activated microglia A” marker genes. Red dotted line indicates −log(p-value) 1.3 (p value 0.05). **d**. Dotplot comparing top 5 marker genes from previous studies of postnatal and disease-associated microglia. (*Clec7a*, not detected in our dataset, was excluded.) **e-g**. RNAscope *in situ* hybridization showing expression of *Spp1* (cyan) and *Gpnmb* (magenta) and *C1qa* (yellow) in uninjured, 1 wpi, and 6 wpi lumbar spinal cord. Scale bars are 200 µm in top row and 50 µm in middle and bottom rows. **h**. Number of Gpnmb+ microglia (quantified by *C1qa, Gpnmb* double-positive cells) in each of the following regions in the spinal cord: dorsal funiculus, dorsal-lateral funiculus, ventral funiculus and grey matter. Error bars indicate mean ± SEM (N = 4 animals). **i-k**. Immunohistochemistry staining for MBP, IBA1 and Neurofilament light, medium and heavy (LMH). Scale bars are 200 µm in top row and 50 µm in bottom row. **l**. Quantification of fluorescence intensity of neurofilament-LMH and MBP in the region boxed i-k, in the dorsal-lateral white matter. * = p-val < 0.001. Error bars indicate mean ± SEM (N = 4 animals).

Notably, the gene expression profile of “activated microglia A” strongly resembled a signature observed recently in postnatal myelin-phagocytosing microglia, in postnatal microglia that can promote SCI repair, and in degenerative disease associated microglia in conditions such as Alzheimer’s disease in the brain and ALS in the spinal cord^27-30^. In addition, recent work examining the lesion site of SCI has identified an “injury associated microglia” cell type with a similar expression profile^31,32^. Pathway analysis of the genes that characterized “activated microglia A” revealed an enrichment of genes associated with (1) phagocytosis (such as *Lyz2, Gpnmb, Itgax*, and *Cd68*; GO terms: lysosome, antigen presentation, phagosome), (2) lipid metabolism (such as *Fabp5, Lgals3, Apoe, Soat1*, and *Abca1*; GO term: lipoprotein), and (3) secreted proteins (such as *Spp1* (OPN) and *Igf1*; GO term “extracellular secretion”, Fig. 4c). We hypothesized that this population, which we term “Trauma Associated Microglia” (TAM) could phagocytose debris in the spared tissue and promote recovery through release of trophic factors.

To determine whether the gene expression pattern that we observed corresponded to an in vivo cell type, we performed in situ hybridization in tissue sections of spared lumbar cords, using C1qa (a general microglial marker), *Spp1* (OPN) (which marked activated microglia as well as some ventral horn neurons, and *Gpnmb* (which marked TAM cells). We found that TAM were observed within the white matter of the injured spinal cord. They appeared consistently along the putative rubrospinal tract (RST) and the dorsolateral corticospinal tract (CST) in the lateral white matter at 1, 3 and 6 wpi (Fig. 4e-h). Interestingly, this region showed loss of longitudinal axons from the descending tracts but also showed the presence of residual myelin (Fig. 4i-l). Myelin debris that remains after longitudinal axons die or retract is thought to be a major obstacle to neuro-regeneration and recovery after SCI^33^.

TAM were also found transiently near the putative main CST at 1 wpi and in the ventral white matter near the descending reticulospinal, vestibulospinal, and cerebellospinal tracts at 6 wpi (Fig. 4f-h). In the cervical spinal cord, rostral to the injury site, TAM were found in a complementary pattern along putative *ascending* tracts (Extended Data Fig. 6). Thus, TAM represented a trauma induced microglial population that was associated with white matter tracts of degenerating axons and that expressed a gene signature of phagocytosis, myelin clearance and lipid processing, and trophic factors.

### *Igf1* and *Spp1* (OPN) Trophic Factors Drive Expansion of TAM Cells

While the phagocytic and lipid processing gene expression modules can be clearly related to a function of TAM, the role and target of the trophic factors *Igf1* and *Spp1* (OPN) in the injured spinal cord is not known. Of note, *Igf1* and *Spp1* (OPN) often function together, with *Spp1* (OPN) sensitizing target cells to *Igf1* signaling^34^. These two secreted proteins are known to be neuroprotective during development^35^ and their co-expression can enable axon regrowth after injury^34,36^. In addition, *Igf1* can prevent oligodendrocyte cell death in transected nerves^37^ and during demyelination^38,39^. Following SCI, we found that TAM were the only cells that co-expressed *Igf1* and *Spp1* (OPN) while many cells, including neurons and oligodendrocytes, expressed the *Igf1r* receptor which would render them responsive to these trophic TAM factors (Fig. 5a).

**Figure 5:**
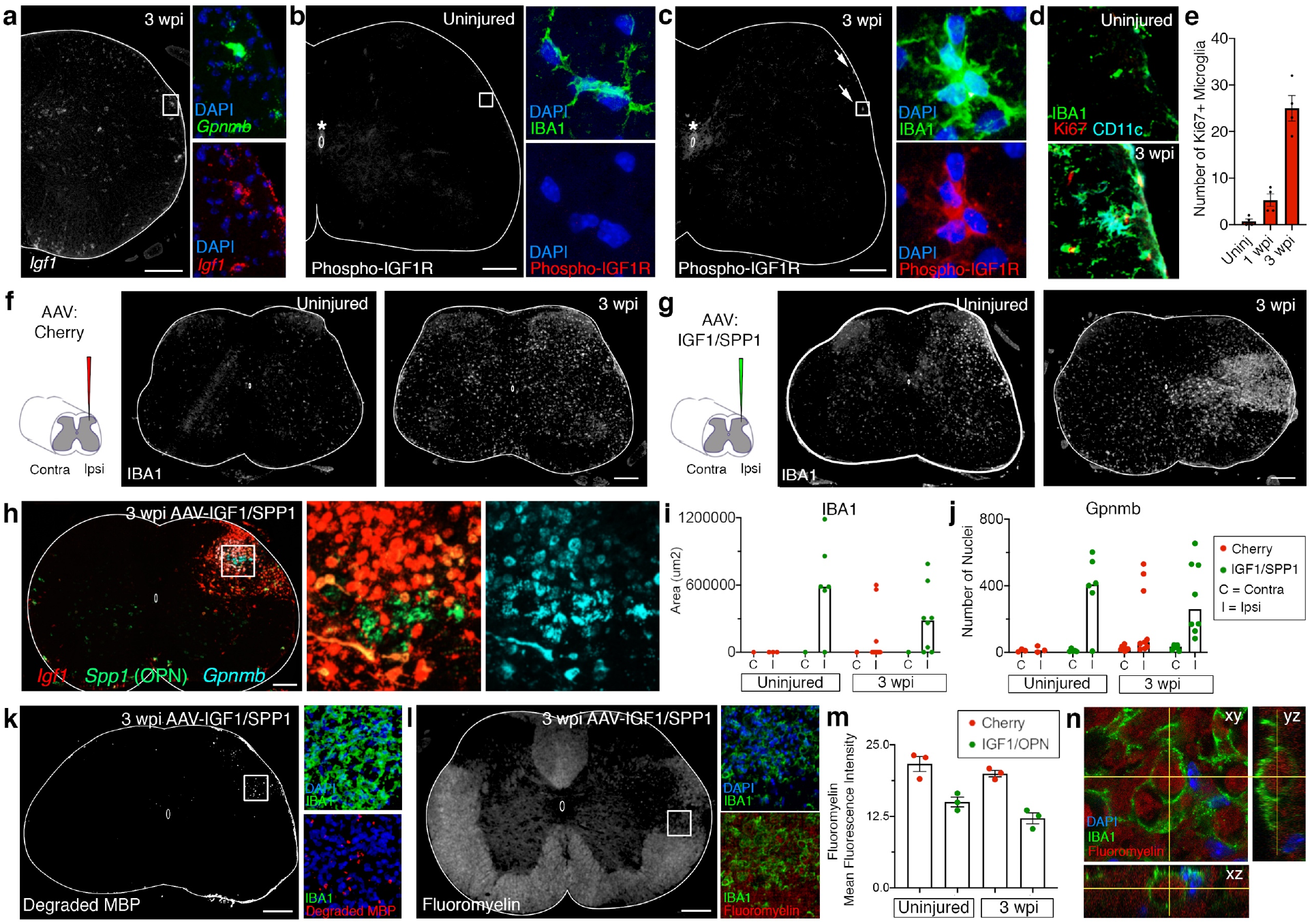
Expression of Trophic Factors Specific to Trauma Associated Microglia Promote Phagocytosis of Myelin. **a**. RNAscope of *Igf1* and *Gpnmb* 3 wpi. **b-c**. Immunohistochemistry of IBA1 and phosphor-IGF1R in the uninjured (**b**) and 3 wpi (**c**) cords. Asterisks represent phospho-IGF1R staining around the central canal. Arrows represent phosphor-IGF1R staining near the dorsal-lateral funiculus. **d**. Immunohistochemistry of IBA1, Ki67 and CD11c in the uninjured and 3 wpi lumbar cord. **e**. Quantification of Ki67+ IBA1+ cells in the uninjured, 1 wpi and 3 wpi cord. **f-g**. Schematic and immunohistochemistry of IBA1 for AAV-cherry **(f)** and AAV-IGF1/SPP1 **(g)** injected cords. Scale bars are 200 µm. **h**. RNAscope *in situ* hybridization of *Igf1* (red), *Spp1* (green), and *Gpnmb* (cyan) at the site of AAV-IGF1/SPP1(OPN) injection. Scale bar is 200 µm. **i**. Quantification of IBA1+ nodule area. Median and individual replicates are shown due to the presence of outliers. **j**. Quantification of the number of Gpnmb+ nuclei. Median and individual replicates are shown due to the presence of outliers. **k**. Immunohistochemistry of degraded MBP and IBA1. **l**. Immunohistochemistry of IBA1 and fluoromyelin. Scale bars are 200 µm. **m**. Quantification of fluoromyelin mean fluorescence intensity after AAV-injection. **n**. Immunohistochemistry of IBA1 and fluoromyelin red, showing fluoromyelin localized within IBA1+ cells at the AAV-IGF1/SPP1(OPN)-injection site 3 wpi. Imaged in multiple confocal sections with xy, xz and yz dimensions shown.

To identify the cellular target of *Igf1* and *Spp1* (OPN) signaling, we analyzed the distribution of the phosphorylated (activated) form of the Igf1 receptor three weeks after thoracic injury, at which point the numbers of TAM cells have increased and plateaued in the lumbar cord. Surprisingly, the only cells which induced high levels of active Igf1r signaling in response to injury were small clusters of microglia that were present in the same white matter spatial distribution as TAM cells, suggesting an autocrine signaling loop (Fig. 5b-c). The small clusters of microglia could represent local proliferation and/or mature cell migration. We analyzed the distribution of the proliferative marker Ki67 together with the microglial marker Iba1 and the TAM protein marker CD11c (*Itgax*). We found that small clusters of microglia in the white matter of the injured spinal cord were indeed proliferating and that some of these cells co-expressed CD11c (Fig. 5d-e). Together, these findings support a model in which TAM cells secrete *Igf1* and *Spp1* (OPN) as autocrine trophic factors to self-amplify following SCI.

To test this hypothesis, we used AAVs to deliver an exogenous source of *Igf1* and *Spp1* (OPN) into the lumbar spinal cord of uninjured mice or following thoracic transection injury and we analyzed the effect on local TAM cells. Three weeks after AAV-injection into the L2 and L5 segments of the lumbar spinal cord, AAV-*Igf1*/*Spp1* (OPN) induced a significant expansion of microglia in both the thoracically-injured and uninjured spinal cords (Fig. 5f-j). These cells formed large nodules of compact cells with ameboid morphology with many examples of active phagocytosis. These microglia expressed *Igf1* and *Spp1* (OPN), as well as the TAM marker *Gpnmb*. Thus, over-expression of *Igf1* and *Spp1* (OPN), which were normally expressed together endogenously in TAM cells, can dramatically expand TAM phagocytic microglia.

Finally, we considered that the ability to increase the TAM population using AAV-*Igf1*/*Spp1* (OPN) may be used to amplify the clearance of residual myelin along degenerating axon tracts. We analyzed the distribution of myelin and of degraded myelin in the context of AAV-*Igf1*/*Spp1* (OPN) expansion of TAM cells and found that, indeed, the region of TAM expansion showed a depletion of myelin (Fig. 5l-n). Moreover, a cryptic epitope of myelin that is only exposed upon myelin degradation was significantly enhanced in response to AAV-*Igf1*/*Spp1* (OPN) (Fig. 5k). Thus, the exogenous manipulation of TAM signaling and population expansion is sufficient to clear myelin debris in spinal cord white matter.

These data demonstrate that SCI induced a small population of TAM in the white matter of the spinal cord, cells with the molecular signature that should enable them to phagocytose and clear myelin. While these TAM cells signal to each other through *Igf1* and *Spp1* (OPN) and proliferate, their natural response is insufficient for removal of myelin debris. However, the exogenous delivery of *Igf1* and *Spp1* (OPN) greatly amplifies this population to facilitate myelin clearance and may provide a pathway for the re-growth of axons that connect the brain with the spinal cord.

## Discussion

SCI disrupts neuronal connections, eliciting trauma on a wide array of cells within the tissue. While the capacity for axonal regeneration and recovery after SCI is limited, ^40,41^ there are latent mechanisms for spontaneous recovery within the spinal cord, particularly when the injury does not sever all connections between the brain and the spinal cord and reorganization of circuits below the lesion can restore function^42-45^. We sought to uncover the endogenous mechanisms by which the adult nervous system *attempts* to recover from an acute SCI, with the hope that this knowledge could inform new therapeutic strategies in the future. In particular, we examined the “spared” lumbar spinal cord after thoracic injury because this tissue has the potential to support recovery in multiple contexts^4-6^. This study includes a single cell atlas that serves as a resource to understand changes in cellular composition, changes in cellular pathways, and specific changes in sub-populations of cells after SCI. From this detailed characterization of cell types, we highlight two unexpected findings that suggested intrinsic attempts at recovery. The first is a rare population of neurons that express a transcriptional signature associated with regeneration, typically seen in regenerating peripheral nerves. Secondly, we identified a population of trauma associated microglia (TAM) distributed along the white matter of degenerating axons and strongly resembled “disease associated microglia” found in Alzheimer’s disease^28,46,47^ and ALS. We found that TAM use autocrine *Igf1*/*Spp1*(OPN) signaling after injury and that exogenous delivery of these factors expanded TAM in vivo and promoted local degradation of myelin debris. In summary, we created a reference atlas of cell-type specific changes after traumatic injury and then used this resource to reveal the small but promising indications of recovery that are present in adult nervous tissue.

This work examined all cell types within the spared tissue below a SCI and complements recent work that has probed specific cellular responses to injury in other locations of the nervous system. This includes the examination of changes in corticospinal neurons in response to SCI^48,49^ as well as astrocytes or myeloid cells at the primary lesion site^31,32,50,51^. Additionally, work profiling the peripheral nervous system has also given important insight into how the nervous system can successfully recovery from injury^15,16^. In comparison with these studies, we also provide a more comprehensive cellular atlas and time course analysis, tracking all cell type changes from 1 dpi through 6 wpi. This broad atlas allowed us to observe both the acute responses to injury and the chronic resolution of many of these changes. At an acute timepoint (1 dpi) we found the greatest change in gene expression across cell types, with many cell types, genes associated with metal binding (*Mt1, Fth1*) were upregulated 1 dpi, suggesting an endogenous protection against the damaging effects of free radicals. In addition, we found pathways associated with structural remodeling across many cell types (Fig. 2b), highlighting subtle cellular adjustments and responses of distal tissues that may be otherwise classified as “normal.”

We next examined a population of neurons that express RAGs after injury, a potential intrinsic attempt at recovery. While CNS neurons have been shown to regenerate after injury^48,49^, RAGs are not typically found in these populations but are rather characteristic of peripheral neuron responses to injury.^15,16,52^ Unexpectedly, we observed that restricted populations of spinal cord neurons express the regenerative signature after contusion or transection injury. The neurons that expressed RAGs were largely excitatory and consisted of several spatially and molecularly distinct populations. Notably, the RAGs are not expressed in all directly injured neurons, as this would encompass lamina I projection neurons but we did not observe RAG expression in this region. While the expression of this signature is highly suggestive of regenerative potential, future work should is required to address the innate possibility for regeneration, utilizing the endogenous expression of RAGs after SCI^13^.

Microglia and hematopoietic cells were the most responsive cell type after injury, beginning in the acute injury setting and persisting in an activated state through chronic timepoints. Of the microglia that respond after SCI, the TAM cells were notable in that they expressed genes similar to proliferative axon tract-associated microglia in postnatal mice^30,53^, disease-associated microglia^28,54^, as well as microglia from the human spinal cord^55^. Furthermore, studies profiling cell types at the lesion site after SCI to have identified similar microglia^31,32^. Based on the widespread presence of a TAM-related cells across injury and disease states, we characterized these cells further.

TAM-related cells co-express two trophic factors, *Igf1* and *Spp1* (OPN) that have known roles in neuroprotection and myelination^46^. Although these factors are required during neuronal development^35^ and enable regeneration of axons after injury^34,36,56,57^, the cellular targets and mechanism by which these actions occur is unknown. Here we provide evidence for how *Igf1* and *Spp1* (OPN) exert their effect on neurons and oligodendrocytes indirectly through their autocrine signaling and expansion of this activated microglial population. Importantly, we found that manipulating this signaling pathway with viral tools allowed TAM to clear the myelin debris that is thought to impede axon regeneration in the central nervous system. This observation is similar to the ability of early postnatal *Spp1*-expressing microglia to remove debris following SCI. Interestingly, these postnatal microglia are activated only briefly, then return to a homeostatic state within a few days following injury which may be a critical feature of their ability to promote wound healing. Relatedly, activated microglia promote OPC differentiation^58^, but it is their subsequent down-regulation or ablation that is important for the successful myelination of regenerated axons^59^. In the future, a temporally regulated delivery of *Igf1* and *Spp1* (OPN) may provide the most effective therapeutic approach to use these factors to remove myelin debris, promote axon regrowth, and remyelination.

There are several limitations to consider in this work. In this study, nuclei were analyzed instead of whole cells because it has been shown that profiling nuclei avoids experimentally-induced gene expression and selective cell death common in single cell profiling^60-62^, though the sequencing depth is lower than for cells^63^. More broadly, while single nucleus sequencing is increasingly transforming our understanding on the biology of disease and trauma, it is important to note that it provides a particular perspective on these complex processes. This approach emphasizes changes observed at the gene expression level and does not address post-transcriptional cellular mechanisms. In addition, it is critical to use independent methods to evaluate the findings from single cell sequencing, as the discrepancy in astrocyte changes between *in silico* and in-tissue data revealed. Despite these limitations, this work provides a powerful reference atlas of cell-type specific changes after traumatic injury and allowed us to discover rare cellular mechanisms directed at tissue repair.

Here, we sought to uncover the endogenous mechanisms by which the adult nervous system *attempts* to recover from an acute SCI. The single cell atlas, discovery of RAG-expressing neurons, and TAM cells each provide important insight into the natural mechanisms of recovery after SCI. Furthermore, the similarity of TAM to recently described populations observed in other injury contexts, Alzheimer’s disease, and ALS suggest shared pathways by which the adult central nervous system responds to trauma. It is hoped that understanding of these intrinsic mechanisms will provide new therapeutic targets to control or even reverse pathological changes across a wide variety of injury and disease.

## Methods

### Mice

All procedures and experiments were approved by the Veterinary Office of the Canton of Geneva (Switzerland) and the National Institute of Neurological Disorders and Stroke Animal Care and Use Committee. Mice for RNA sequencing were female C57BL/6 (12-30 weeks of age). For all other experiments, balanced samples of male and female C57BL/6 mice (12-30 weeks of age) were used.

### Surgical Procedures

Surgical procedures were performed as previously described^64^. Briefly, following a mid-thoracic laminectomy (T9 vertebra), a spinal cord impactor (IH-0400 Impactor, Precision Systems and Instrumentation LLC) was used to induce a contusion injury. The applied force was set to 90 kdyn. Spinal transections were performed following a mid-thoracic laminectomy (T9 vertebra), by cutting of the spinal cord with spring scissors, before filling the void with gel-foam. Animal care, including manual bladder voiding, was performed twice daily or as needed following injury.

### Viruses

AAV2/1-hSyn mCherry virus was produced at Vigene Biosciences. AAV2/1-IGF1^34^, AAV2/1-OPN^57^ and AAV2/1-PLAP^65^ viruses were a gift from Zhigang He and produced at Boston Children’s Hospital. Viral particles were injected at a titer of 5^E12^-1^E13^ genome copies per ml.

### Intraspinal Injections

Intraspinal injections were performed as previously described^66^. Briefly, mice were anesthetized by intraperitoneal injection of a drug cocktail containing fentanyl (0.2 mg.kg), dexmedetomidin (1 mg/kg), and midazolam (5 mg/kg) dissolved in saline. For spinal injections, a small incision was made in the skin and the underlying musculature and adipose tissue was teased apart to reveal the vertebral column. Tissue joining the dorsal processes of consecutive vertebrae was removed and the vertebral surfaces were cleaned with fine forceps and gently separated to reveal the dorsal surface of the spinal cord (at spinal levels L2 and L5). The dura was punctured by pinching with sharp forceps to facilitate smooth entry of the needle. Virus was pressure injected through a pulled glass needle at a depth of 250 μm from the dorsal surface and 250 nL of viral particles was released at a rate of 100 nl/min.

Following injections, the overlying muscle was sutured, and the skin incision was closed using wound clip. Anesthesia was reversed by intraperitoneal administration of buprenorphine (0.1 mg/kg), atipemazole (2.5 mg/kg), and flumenazil (0.2 mg/kg) in saline. Additionally, mice received intradermal injection of meloxicam SR for analgesia and were returned to their home cages.

### Behavioral assessments

All procedures have been described in detail previously^64^. Limb motor movements was evaluated while running on a horizontal walkway. Bilateral leg kinematics were captured with the Vicon Motion Systems, UK (combining 12 infrared cameras) for tracking with reflective markers on the crest, hip, knee, ankle joints and distal toes. The limbs were modelled as an interconnected chain of segments and gait parameters were calculated from those.

### Analysis of kinematic data

A total of 78 gait parameters were computed for each limb for each gait cycle. We chose to represent the following gait parameters: step height, drag percentage, amp limb, amp speed limb, amp join 1, amp joint 2, amp joint 3, and amp speed joint 3. Differences among groups were calculated using two-tailed t tests (unpaired) and were considered significant if p < 0.05. Data are represented as mean ± SEM unless otherwise indicated. Statistical analyses were performed using GraphPad prism software.

### Animal Inclusion and Exclusion Criteria

Two days after injury, all mice were evaluated in an open field and all animals exhibiting any hindlimb movements were not further studied. A larger cohort of mice were taken through kinematic analysis, and 3 mice representative of each timepoint were selected for snRNAseq.

### Nuclei Isolation

Nuclei were isolated from adult mouse lumbar cords with proximal dorsal and ventral roots attached using a triton-based protocol adapted from Matson et al. 2018. Briefly, mice were euthanized according to IACUC guidelines. The spinal cord was rapidly dissected and frozen on dry ice. Later, fresh frozen lumbar cords (spinal segment L2-S1) were placed in a Dounce homogenizer (Kontes Dounce Tissue Grinder) containing 500 μL of lysis buffer (0.32 M sucrose, 10 mM HEPES [pH 8.0], 5 mM CaCl_2_, 3 mM MgAc, 0.1 mM ETDA, 1 mM DTT, 0.1% Triton X-100). The cords were dounced with 5 strokes of pestle A, then 5-10 strokes of pestle B. The lysate was diluted in 3 mL of sucrose buffer (0.32 M sucrose, 10 mM HEPES [pH 8.0], 5 mM CaCl_2_, 3 mM MgAc, 0.1 mM ETDA, 1 mM DTT) and passed over a 40 μm strainer. The filtered lysate was centrifuged at 3,200 x g for 10 min at 4°C. After centrifugation, the pellet was resuspended in sucrose buffer and incubated for 2 min on ice. The sample was transferred to an Oak Ridge tube and homogenized for 1 min using an Ultra-Turrax Homogenizer (IKA). Then, 12.5 mL of density sucrose buffer (1 M sucrose, 10 mM HEPES [pH 8.0], 3 mM MgAc, 1 mM DTT) was layered below the sample. The tube was centrifuged at 3,200 x g for 20 min and the supernatant immediately poured off. The nuclei on the side of the tube were resuspended with 100 μL of PBS with 0.04% BSA and 0.2 U/ μL RNase inhibitor. Nuclei were inspected for visual appearance and quantified with a hemocytometer before being adjusted to a concentration of 1,000 nuclei per μL.

### Single Nucleus RNA Sequencing

Single nucleus RNA sequencing was carried out using Single-cell gene expression 3’ v2 kit on the Chromium platform (10X Genomics) according to manufacturer’s instructions with one modification. Following reverse-transcription, an additional PCR cycle was added to the number of cycles for cDNA amplification to compensate for decreased cDNA abundance in nuclei compared to cells. Approximately 8,000-9,000 nuclei were loaded in each well and 3,000-7,000 nuclei were recovered per sample.

Libraries were sequenced to a minimum depth of 20,000 reads per nucleus using an Illumina HiSeq 3000 (PE 26 – 8 – 98 bp). Raw sequencing reads were demultiplexed, aligned, and a count matrix was generated using CellRanger. For alignment, introns and exons were included in the reference genome (mm10).

### Clustering

Seurat v3.2.2 was used to filter, normalize, anchor, and cluster the dataset^12^. We filtered nuclei for downstream analysis, including those with greater than 200 genes per nucleus and less than 20 percent of reads coming from mitochondrial genes. For neurons, the minimum threshold was increased to 500 genes per nucleus. All genes analyzed were present in greater than three nuclei. We performed standard log normalization and used principal component analysis for dimensionality reduction. We used the FindIntegrationAnchors and FindTransferAnchors functions to set the uninjured samples as reference anchors for coarse clustering. For the re-clustering of microglia/hematopoietic cell, given the increase in diversity of cell types and increase in number after injury, FindIntegration and FindTransfer were run without using uninjured samples as a reference anchor. Clusters were visualized using Uniform Manifold Approximation and Projection for Dimension Reduction (UMAP), and cluster markers were found using the “auroc” test in Seurat. Clusters with less than 3 significant markers, or that were not defined by a cohesive set of genes, were identified as low-quality clusters and discarded from downstream analysis. Clusters that contained nuclei that expressed more than 3 genes from two cell types were identified as doublets and removed from downstream analysis. Nuclei had on average 1392 genes per nucleus in neurons and 471 genes per nucleus in non-neuronal cells. Neurons were classified using label transfer^12^ at a “family”-level resolution^11^.

### Pathway and Cell Type Prioritization

Augur was implemented as previously described.^9^ For pathway analysis, differential gene expression across conditions was generated using FindMarkers using the Wilcox test. GO Analysis was done using all differentially expressed genes with p_adj < 0.05, GO BP, CC, MF -> clustering. In cases in which multiple clusters had the same genes and similar terms, only the most significant is shown.

### Immunohistochemistry Antibodies

IBA1 (Cedarlane Labs, 234006(SY)), TMEM 119 (Cedarlane Labs, 400004(SY)), CD11c (GeneTex, GTX74935), Myelin-MBP (BioLegend, 808402), NF-L (Cell Signaling, 2835S), NF-M (Cell Signaling, 2838S), NF-H (Cell Signaling, 2836S), NeuN (Millipore Sigma, ABN90P), CD68 (Abcam, ab125212), CNPase (Millipore Sigma, MAB326), GFAP (Agilent/Dako, Z033429-2), Lectin (Vector Labs, DL-1177-1), DAPI, Cleaved Caspase 3 (Cell Signalling Tech, 9661L), Cleaved PARP (Cell Signalling Tech, 94885S), Phospho-IGF1R (Invitrogen, PA5-104773), Fluoromyelin Red (Invitrogen, F34652), Degraded MBP (Sigma-Aldrich, AB5864).

### *In Situ* Hybridization Probes

Spp1 (435191), Vsx2 (438341), Gap43 (318621), Chat (408731), Itgax (311501), Mdga1 (546411), Sprr1a (426871-C2), Vgf (517421-C2), Igf1 (443901-C2), Sprr1a (426871-C2), C1qa (441221-C2), Megf11 (504281-C2), GFP (409011-C2), Apoe (313271-C3), Tnfrsf12a (429311-C3), Atf3 (426891-C3), Gpr83 (317431-C3), Pdgfra (480661-C3), Shox2 (554291-C3), Gpnmb (489511-C3).

### Immunohistochemistry and *In Situ* Hybridization

Animals were euthanized with avertin and perfused with PBS and then 4% paraformaldehyde. Spinal cords were dissected, fixed in 4% paraformaldehyde overnight, washed in PBS for one hour, then dehydrated in 30% sucrose an additional night before being embedded in OCT.

Immunohistochemistry was performed as previously described^61^. Briefly, spinal cords were cut at 50 µm, placed in blocking buffer (1% IgG-free BSA, 10% normal donkey serum, 0.1% Triton-X 100 in PBS) for one hour, prior to incubation in blocking buffer and primary antibody for 48 hours at 4°C. Primary antibody was washed off three times in PBS before a 2-hour incubation in secondary antibody at room temperature. Secondary antibody was washed off three times in PBS before adding a coverslip.

Multiplex immunohistochemistry was performed as previously described^8^.

*In situ* hybridization was performed according to manufacturer’s instructions for fixed frozen tissue RNAscope (ACD Bio).

### Imaging

Images of immunohistochemistry and *in situ* hybridization samples were imaged using a Zeiss 800 LSM confocal microscope. For quantification, a tile scan image spanning the section was generated for ≥ 3 sections from ≥ 3 mice. Brightness and contrast were adjusted in Photoshop (Adobe), standardized across images.

### Quantification of Cell Counts from Images of Immunohistochemistry

A custom MATLAB-based image analysis program was developed to assist the cell counting for this work. The software automatically identifies and counts cells based on a criterion that constrains size at a user-selectable intensity threshold. A manual selection tool is also available to identify additional cells that are more difficult to detect. A second channel of the same image shows can be used to counts cells that are labeled with both stains using the results from the first channel and a second set of user-selectable thresholds.

### Fluorescence Intensity

Mean fluorescence intensity was quantified using FIJI (ImageJ).

### Histological Quantification and Statistical Testing

Two-tailed t tests (unpaired) were used for quantification of immunohistochemistry and *in situ* hybridization. Differences among groups were considered significant if p < 0.05. Data are represented as mean ± SEM unless otherwise indicated. Statistical analyses were performed using GraphPad prism software.

## Data Availability

A searchable version of this data is available at https://seqseek.ninds.nih.gov/spinalcordinjury. Raw sequencing data and count matrices have been deposited to the Gene Expression Omnibus. Custom MATLAB-based code for quantification of cell counts is available at https://github.com/ArielLevineLabNINDS/CellCounter.

## Acknowledgements

The authors gratefully acknowledge technical support from Li Li, Aaron Bickert and Stefan Stoica, and advice and comments from Claire Le Pichon and Michael O’Donovan, as well as the veterinary staff of the Porter Research Building, including Heather Narver. We thank Yuanyuan Liu (NIH, NIDCR) and Zhigang He (Boston Children’s Hospital) for the AAV-IGF/SPP1(OPN) viruses. This research was supported in part by the Intramural Research Program of the NIH, NINDS, and CIT. A.S., I.H., K.J.E.M., and A.J.L. were supported by NIH Intramural Funds through 1ZIA NS003153; and J.C. and R.P. were supported by Z99 CT999999. G.C., C.K. and J.W.S. were supported by the European Research Council [ERC-2015-CoG HOW2WALKAGAIN 682999] and the Swiss National Science Foundation (subside 310030_192558).

## Author Contributions

This project was conceptualized by A.J.L. and K.J.E.M with input from C.K., J.W.S. and G.C. Experiments were performed by K.J.E.M., D.M. and C.K. Data was analyzed by K.J.E.M., D.E.R., J.W.S., J.C., R.P., I.H., and A.S. The manuscript was written by K.J.E.M. and A.J.L with input from all authors. Supervision and funding acquisition were by A.J.L. and G.C.

## References

1 Kigerl, K. A. et al. Identification of two distinct macrophage subsets with divergent effects causing either neurotoxicity or regeneration in the injured mouse spinal cord. J Neurosci 29, 13435–13444, doi:10.1523/JNEUROSCI.3257-09.2009 (2009).

2 Sofroniew, M. V. & Vinters, H. V. Astrocytes: biology and pathology. Acta Neuropathol 119, 7–35, doi:10.1007/s00401-009-0619-8 (2010).

3 McDonald, J. W. & Sadowsky, C. Spinal-cord injury. Lancet 359, 417–425, doi:10.1016/S0140-6736(02)07603-1 (2002).

4 Angeli, C. A. et al. Recovery of Over-Ground Walking after Chronic Motor Complete Spinal Cord Injury. N Engl J Med 379, 1244–1250, doi:10.1056/NEJMoa1803588 (2018).

5 Gill, M. L. et al. Neuromodulation of lumbosacral spinal networks enables independent stepping after complete paraplegia. Nat Med 24, 1677–1682, doi:10.1038/s41591-018-0175-7 (2018).

6 Wagner, F. B. et al. Targeted neurotechnology restores walking in humans with spinal cord injury. Nature 563, 65–71, doi:10.1038/s41586-018-0649-2 (2018).

7 Matson, K. J. E. et al. Isolation of Adult Spinal Cord Nuclei for Massively Parallel Single-nucleus RNA Sequencing. J Vis Exp, doi:10.3791/58413 (2018).

8 Maric, D. et al. Whole-brain tissue mapping toolkit using large-scale highly multiplexed immunofluorescence imaging and deep neural networks. Nat Commun 12, 1550, doi:10.1038/s41467-021-21735-x (2021).

9 Skinnider, M. A. et al. Cell type prioritization in single-cell data. Nat Biotechnol 39, 30–34, doi:10.1038/s41587-020-0605-1 (2021).

10 Liddelow, S. A. et al. Neurotoxic reactive astrocytes are induced by activated microglia. Nature 541, 481–487, doi:10.1038/nature21029 (2017).

11 Russ, D. E., Patterson Cross, R. B., Li, L., Koch, S. C., Matson, K. J. E., Levine, A. J.. A Harmonized Atlas of Spinal Cord Cell Types and Their Computational Classification. bioRxiv, doi:doi: https://doi.org/10.1101/2020.09.03.241760 (2020).

12 Stuart, T. et al. Comprehensive Integration of Single-Cell Data. Cell 177, 1888–1902 e1821, doi:10.1016/j.cell.2019.05.031 (2019).

13 Holland, S. D., Ramer, L. M., McMahon, S. B., Denk, F. & Ramer, M. S. An ATF3-CreERT2 Knock-In Mouse for Axotomy-Induced Genetic Editing: Proof of Principle. eNeuro 6, doi:10.1523/ENEURO.0025-19.2019 (2019).

14 He, Z. & Jin, Y. Intrinsic Control of Axon Regeneration. Neuron 90, 437–451, doi:10.1016/j.neuron.2016.04.022 (2016).

15 Renthal, W. et al. Transcriptional Reprogramming of Distinct Peripheral Sensory Neuron Subtypes after Axonal Injury. Neuron 108, 128–144 e129, doi:10.1016/j.neuron.2020.07.026 (2020).

16 Nguyen, M. Q., Le Pichon, C. E. & Ryba, N. Stereotyped transcriptomic transformation of somatosensory neurons in response to injury. Elife 8, doi:10.7554/eLife.49679 (2019).

17 Barik, A. et al. A spinoparabrachial circuit defined by Tacr1 expression drives pain. Elife 10, doi:10.7554/eLife.61135 (2021).

18 Haring, M. et al. Neuronal atlas of the dorsal horn defines its architecture and links sensory input to transcriptional cell types. Nat Neurosci 21, 869–880, doi:10.1038/s41593-018-0141-1 (2018).

19 Choi, S. et al. Parallel ascending spinal pathways for affective touch and pain. Nature 587, 258–263, doi:10.1038/s41586-020-2860-1 (2020).

20 Roome, R. B. et al. Phox2a Defines a Developmental Origin of the Anterolateral System in Mice and Humans. Cell Rep 33, 108425, doi:10.1016/j.celrep.2020.108425 (2020).

21 Osseward, P. J., 2nd et al. Conserved genetic signatures parcellate cardinal spinal neuron classes into local and projection subsets. Science 372, 385–393, doi:10.1126/science.abe0690 (2021).

22 Hayashi, M. et al. Graded Arrays of Spinal and Supraspinal V2a Interneuron Subtypes Underlie Forelimb and Hindlimb Motor Control. Neuron 97, 869–884 e865, doi:10.1016/j.neuron.2018.01.023 (2018).

23 Delile, J. et al. Single cell transcriptomics reveals spatial and temporal dynamics of gene expression in the developing mouse spinal cord. Development 146, doi:10.1242/dev.173807 (2019).

24 Dougherty, K. J. et al. Locomotor rhythm generation linked to the output of spinal shox2 excitatory interneurons. Neuron 80, 920–933, doi:10.1016/j.neuron.2013.08.015 (2013).

25 Ha, N. T. & Dougherty, K. J. Spinal Shox2 interneuron interconnectivity related to function and development. Elife 7, doi:10.7554/eLife.42519 (2018).

26 Baek, M., Menon, V., Jessell, T. M., Hantman, A. W. & Dasen, J. S. Molecular Logic of Spinocerebellar Tract Neuron Diversity and Connectivity. Cell Rep 27, 2620–2635 e2624, doi:10.1016/j.celrep.2019.04.113 (2019).

27 Hammond, T. R. et al. Single-Cell RNA Sequencing of Microglia throughout the Mouse Lifespan and in the Injured Brain Reveals Complex Cell-State Changes. Immunity 50, 253–271 e256, doi:10.1016/j.immuni.2018.11.004 (2019).

28 Keren-Shaul, H. et al. A Unique Microglia Type Associated with Restricting Development of Alzheimer’s Disease. Cell 169, 1276–1290 e1217, doi:10.1016/j.cell.2017.05.018 (2017).

29 Li, Y. et al. Microglia-organized scar-free spinal cord repair in neonatal mice. Nature 587, 613–618, doi:10.1038/s41586-020-2795-6 (2020).

30 Li, Q. et al. Developmental Heterogeneity of Microglia and Brain Myeloid Cells Revealed by Deep Single-Cell RNA Sequencing. Neuron 101, 207–223 e210, doi:10.1016/j.neuron.2018.12.006 (2019).

31 Wahane, S. et al. Diversified transcriptional responses of myeloid and glial cells in spinal cord injury shaped by HDAC3 activity. Sci Adv 7, doi:10.1126/sciadv.abd8811 (2021).

32 Millich, L. M., Choi, J., Ryan, C., Yahn, S.L., Tsoulfas, P., Lee, J.K. Single cell analysis of the cellular heterogeneity and interactions in the injured mouse spinal cord. bioRxiv, doi:https://doi.org/10.1101/2020.05.13.094854 (2020).

33 Schwab, M. E. & Strittmatter, S. M. Nogo limits neural plasticity and recovery from injury. Curr Opin Neurobiol 27, 53–60, doi:10.1016/j.conb.2014.02.011 (2014).

34 Liu, Y. et al. A Sensitized IGF1 Treatment Restores Corticospinal Axon-Dependent Functions. Neuron 95, 817–833 e814, doi:10.1016/j.neuron.2017.07.037 (2017).

35 Ueno, M. et al. Layer V cortical neurons require microglial support for survival during postnatal development. Nat Neurosci 16, 543–551, doi:10.1038/nn.3358 (2013).

36 Anderson, M. A. et al. Required growth facilitators propel axon regeneration across complete spinal cord injury. Nature 561, 396–400, doi:10.1038/s41586-018-0467-6 (2018).

37 Barres, B. A., Jacobson, M. D., Schmid, R., Sendtner, M. & Raff, M. C. Does oligodendrocyte survival depend on axons? Curr Biol 3, 489–497, doi:10.1016/0960-9822(93)90039-q (1993).

38 Mason, J. L. et al. Mature oligodendrocyte apoptosis precedes IGF-1 production and oligodendrocyte progenitor accumulation and differentiation during demyelination/remyelination. J Neurosci Res 61, 251–262, doi:10.1002/1097-4547(20000801)61:3<251::AID-JNR3>3.0.CO;2-W (2000).

39 Mason, J. L., Ye, P., Suzuki, K., D’Ercole, A. J. & Matsushima, G. K. Insulin-like growth factor-1 inhibits mature oligodendrocyte apoptosis during primary demyelination. J Neurosci 20, 5703–5708 (2000).

40 O’Shea, T. M., Burda, J. E. & Sofroniew, M. V. Cell biology of spinal cord injury and repair. J Clin Invest 127, 3259–3270, doi:10.1172/JCI90608 (2017).

41 Tran, A. P., Warren, P. M. & Silver, J. The Biology of Regeneration Failure and Success After Spinal Cord Injury. Physiol Rev 98, 881–917, doi:10.1152/physrev.00017.2017 (2018).

42 Courtine, G. et al. Recovery of supraspinal control of stepping via indirect propriospinal relay connections after spinal cord injury. Nat Med 14, 69–74, doi:10.1038/nm1682 (2008).

43 Hilton, B. J. & Bradke, F. Can injured adult CNS axons regenerate by recapitulating development? Development 144, 3417–3429, doi:10.1242/dev.148312 (2017).

44 van den Brand, R. et al. Restoring voluntary control of locomotion after paralyzing spinal cord injury. Science 336, 1182–1185, doi:10.1126/science.1217416 (2012).

45 Murray, K. C. et al. Recovery of motoneuron and locomotor function after spinal cord injury depends on constitutive activity in 5-HT2C receptors. Nat Med 16, 694–700, doi:10.1038/nm.2160 (2010).

46 Benmamar-Badel, A., Owens, T. & Wlodarczyk, A. Protective Microglial Subset in Development, Aging, and Disease: Lessons From Transcriptomic Studies. Front Immunol 11, 430, doi:10.3389/fimmu.2020.00430 (2020).

47 Kamphuis, W., Kooijman, L., Schetters, S., Orre, M. & Hol, E. M. Transcriptional profiling of CD11c-positive microglia accumulating around amyloid plaques in a mouse model for Alzheimer’s disease. Biochim Biophys Acta 1862, 1847–1860, doi:10.1016/j.bbadis.2016.07.007 (2016).

48 Fink, K. L., Lopez-Giraldez, F., Kim, I. J., Strittmatter, S. M. & Cafferty, W. B. J. Identification of Intrinsic Axon Growth Modulators for Intact CNS Neurons after Injury. Cell Rep 18, 2687–2701, doi:10.1016/j.celrep.2017.02.058 (2017).

49 Poplawski, G. H. D. et al. Injured adult neurons regress to an embryonic transcriptional growth state. Nature 581, 77–82, doi:10.1038/s41586-020-2200-5 (2020).

50 Anderson, M. A. et al. Astrocyte scar formation aids central nervous system axon regeneration. Nature 532, 195–200, doi:10.1038/nature17623 (2016).

51 Chen, K. et al. RNA-seq characterization of spinal cord injury transcriptome in acute/subacute phases: a resource for understanding the pathology at the systems level. PLoS One 8, e72567, doi:10.1371/journal.pone.0072567 (2013).

52 Starkey, M. L. et al. Expression of the regeneration-associated protein SPRR1A in primary sensory neurons and spinal cord of the adult mouse following peripheral and central injury. J Comp Neurol 513, 51–68, doi:10.1002/cne.21944 (2009).

53 Hammond, T. R., Robinton, D. & Stevens, B. Microglia and the Brain: Complementary Partners in Development and Disease. Annu Rev Cell Dev Biol 34, 523–544, doi:10.1146/annurev-cellbio-100616-060509 (2018).

54 Krasemann, S. et al. The TREM2-APOE Pathway Drives the Transcriptional Phenotype of Dysfunctional Microglia in Neurodegenerative Diseases. Immunity 47, 566–581 e569, doi:10.1016/j.immuni.2017.08.008 (2017).

55 Shannon Tansley, S. U., Alba Ureña Guzmán, Moein Yaqubi, Alain Pacis, Marc Parisien, Oded Rabau, Lisbet Haglund, Jean Ouellet, Carlo Santaguida, Jiannis Ragoussis, Ji Zhang, Michael W Salter, Luda Diatchenko, Luke M. Healy, Jeffrey S. Mogil, Arkady Khoutorsky. Single-cell RNA sequencing reveals time- and sex-specific responses of spinal cord microglia to peripheral nerve injury and links ApoE to neuropathic pain. bioRxiv, doi:https://doi.org/10.1101/2020.12.09.418541 (2020).

56 Bei, F. et al. Restoration of Visual Function by Enhancing Conduction in Regenerated Axons. Cell 164, 219–232, doi:10.1016/j.cell.2015.11.036 (2016).

57 Duan, X. et al. Subtype-specific regeneration of retinal ganglion cells following axotomy: effects of osteopontin and mTOR signaling. Neuron 85, 1244–1256, doi:10.1016/j.neuron.2015.02.017 (2015).

58 Lloyd, A. F. & Miron, V. E. The pro-remyelination properties of microglia in the central nervous system. Nat Rev Neurol 15, 447–458, doi:10.1038/s41582-019-0184-2 (2019).

59 Wang, J. et al. Robust Myelination of Regenerated Axons Induced by Combined Manipulations of GPR17 and Microglia. Neuron 108, 876–886 e874, doi:10.1016/j.neuron.2020.09.016 (2020).

60 Lacar, B. et al. Nuclear RNA-seq of single neurons reveals molecular signatures of activation. Nat Commun 7, 11022, doi:10.1038/ncomms11022 (2016).

61 Sathyamurthy, A. et al. Massively Parallel Single Nucleus Transcriptional Profiling Defines Spinal Cord Neurons and Their Activity during Behavior. Cell Rep 22, 2216–2225, doi:10.1016/j.celrep.2018.02.003 (2018).

62 Alkaslasi, M. R. P. Z.E., Silberberg, H., Chen, L., Zhang, Y., Petros, T.J., Le Pichon, C.E. Single nucleus RNA-sequencing defines unexpected diversity of cholinergic neuron types in the adult mouse spinal cord. bioRxiv, doi:https://doi.org/10.1101/2020.07.16.193292 (2021).

63 Bakken, T. E. et al. Single-nucleus and single-cell transcriptomes compared in matched cortical cell types. PLoS One 13, e0209648, doi:10.1371/journal.pone.0209648 (2018).

64 Asboth, L. et al. Cortico-reticulo-spinal circuit reorganization enables functional recovery after severe spinal cord contusion. Nat Neurosci 21, 576–588, doi:10.1038/s41593-018-0093-5 (2018).

65 Liu, K. et al. PTEN deletion enhances the regenerative ability of adult corticospinal neurons. Nat Neurosci 13, 1075–1081, doi:10.1038/nn.2603 (2010).

66 Sathyamurthy, A. et al. Cerebellospinal Neurons Regulate Motor Performance and Motor Learning. Cell Rep 31, 107595, doi:10.1016/j.celrep.2020.107595 (2020).

